# The selective OX1R antagonist 1-SORA-51 reduces binge-like feeding behavior in male and female mice without detectable changes in dopamine

**DOI:** 10.64898/2026.04.24.720455

**Authors:** J.R. Trinko, E. Atangana, D. Diaz, A. Ashkenazi, E. Foscue, E. Kong, R. J. DiLeone

## Abstract

Binge eating disorder (BED) is characterized by episodic overconsumption of palatable food and involves dysregulated motivational and arousal processes. The orexin (hypocretin) system, through its widespread projections to mesolimbic circuits, has been implicated in cue-driven reward seeking and escalated intake, raising the possibility that orexin receptor antagonists may modulate binge-like behavior. Here we evaluated the effects of a dual orexin receptor antagonist (DORA-22) and an OX1R-selective antagonist (1-SORA-51) in a cyclic intermittent high-fat access model that generates robust and reproducible binge-like intake in male and female mice. DORA-22 produced no detectable effect on consumption at either early (2 h) or extended (24 h) binge timepoints. In contrast, 1-SORA-51 significantly reduced high-fat intake during the initial 2 hours of access in both sexes, with no effect on 24-hour consumption, indicating a selective attenuation of the early phase of binge intake. Fiber photometry recordings of GRABDA_2m_ fluorescence in the nucleus accumbens revealed that 1-SORA-51 did not alter baseline dopamine signals or the dopamine increase triggered by high-fat pellet delivery, demonstrating that its behavioral effects occur without detectable modulation of mesolimbic dopamine dynamics. Together, these findings identify OX1R antagonism as a strategy for suppressing the initiation of binge-like feeding and highlight the receptor-level specificity of orexin contributions to maladaptive overconsumption.

## Introduction

Binge eating disorder (BED) is defined by the Diagnostic and Statistical Manual of Mental Disorders (DSM-5) as an eating disorder that combines aspects of overeating with negative psychological or social effects (*Diagnostic and Statistical Manual of Mental Disorders, Fifth Edition* 2013). Physical features of BED include unusually rapid consumption of large quantities of food, often when not hungry and to the point of discomfort. Psychologically, episodes are marked by a sense of loss of control and followed by pronounced negative affect, including guilt, shame, or disgust, often in the context of disturbed body image. Recurrent episodes occur without compensatory behavior such as purging. BED is the most prevalent eating disorder, has high comorbidities with obesity, metabolic syndrome, and psychiatric disorders (Hudson et al. 2007; Swanson et al. 2011; Udo and Grilo 2018), and disproportionately affects females (Hudson et al. 2007).

Binge episodes arise from dysregulated interactions among the neural systems that govern energy balance, reward valuation, cue responsivity, cognitive control and affect; contemporary models emphasize that these domains form an integrated control architecture rather than separable homeostatic and hedonic modules (Rossi and Stuber 2018; Hardaway et al. 2015). Within this framework, the tendency of individuals with BED to consume highly palatable, energy-dense foods resembles compulsive intake patterns characteristic of substance use disorders (Volkow et al. 2013; Corwin and Buda-Levin 2004). Consistent with this, reward-circuit abnormalities are frequently reported in BED, with individuals showing altered striatal dopamine responsivity to palatable food cues. However, the direction of dysregulation (hypo-versus hyperdopaminergic) varies across studies, likely owing to differences in task structure, metabolic state, and disease severity. Despite this heterogeneity, converging human and preclinical data support the involvement of mesolimbic dopamine pathways in binge-like feeding, underscoring the relevance of neuromodulatory systems, including orexin, that interface with these circuits (Tomasi and Volkow 2013; Yu et al. 2022; Wang et al. 2011).

Orexins (also termed hypocretins) are peptide neurotransmitters produced exclusively by neurons in the lateral hypothalamus and project widely to target neurons expressing the orexin receptors OX1R and OX2R (Sakurai et al. 1998; de Lecea et al. 1998). Orexin neurons were first implicated in feeding behavior when early studies reported that fasting increased orexin expression (Cai et al. 1999), an observation that initially positioned the peptide as a potential coordinator of responses to energetic challenge. As subsequent studies in sleep and arousal physiology clarified, however, the dominant and evolutionarily conserved function of orexin signaling is to stabilize wakefulness and sustain goal-directed engagement. The discovery that loss of orexin signaling causes narcolepsy (Lin et al. 1999) established the system as a master regulator of arousal state and ultimately enabled the development of dual orexin receptor antagonists such as suvorexant for treatment of insomnia (Winrow et al. 2011; Herring et al. 2016). Within this broader framework, orexin-related changes in feeding are now viewed as downstream consequences of its role in mobilizing attention, locomotion, and behavioral activation.

Orexin’s capacity to amplify behavioral activation makes it a potent modulator of neural circuits governing motivation and reward. Orexin neurons project densely to mesolimbic structures, including the ventral tegmental area and ventral striatum, where orexin receptor signaling enhances dopaminergic responsiveness to salient cues and facilitates reward seeking (Thorpe and Kotz 2005; Carr 2002; Cluderay et al. 2002; Levin 2006; Marcus et al. 2001; Brown et al. 2016; Lai et al. 2018). Through these pathways, orexin contributes to the vigor and persistence of pursuit behaviors across domains, from drug self-administration to palatable food intake, positioning the system as a key amplifier of compulsive or escalated intake. This intersection of arousal-state regulation with mesolimbic modulation has motivated growing interest in targeting orexin receptors to attenuate maladaptive consumption patterns, including those underlying binge-like eating.

Pharmacological suppression of orexin signaling has been evaluated across multiple rodent paradigms of escalated or binge-like intake. Dual orexin receptor antagonists such as SB-649868 reduce high-fat diet consumption in rats (Piccoli et al. 2012), whereas the OX1R-selective antagonist SB-334867 decreases sucrose- or saccharin-driven binge-like drinking in mice (Alcaraz-Iborra et al. 2014). Interpretation across studies, however, is complicated by substantial methodological heterogeneity, with models varying in the degree to which they incorporate stressors (e.g., food restriction, footshock), limited-access timing, “drinking-in-the-dark” procedures, or intermittent presentation of highly palatable foods (Rehn et al. 2025; Cowin et al. 2011). To isolate binge-like intake driven primarily by opportunity and palatability rather than stress or deprivation, we employed the cyclic intermittent high-fat access paradigm developed by Czyzyk et al. (2010) and used this model to examine the effects of the dual antagonist DORA-22 and the OX1R-selective antagonist 1-SORA-51 on feeding behavior and nucleus accumbens dopamine dynamics (Czyzyk et al. 2010).

## Results

### Intermittent access to a palatable diet induces binge-like feeding behavior

To confirm that the intermittent high-fat exposure paradigm reliably elicits binge-like intake within the early time window relevant for pharmacological testing, we first replicated the Czyzyk et al. (2010) protocol in male and female mice. Male and female mice were exposed to the palatable high-fat diet over 48 hours, along with standard chow. Following this pre-exposure, mice were provided with both the high-fat diet and chow, for 24-hour binge sessions, each separated by 6 days of chow alone. As expected, both male and female mice displayed significantly increased food consumption during the binge sessions at 24 hours compared to their chow only consumption the previous day (**Fig 1A**, males, paired t-test, n=10, Binge 1: t=18.49, \*\*\*\**p*<0.0001, Binge 2: t=15.57, \*\*\*\**p*<0.0001; **Fig 1B**, females paired t-test, n=10, Binge 1: t=6.422, \*\*\**p*=0.0001, Binge 2: t=15.21, \*\*\*\**p*<0.0001). Importantly, this escalation was evident in the first 2 hours, with both sexes showing significant increases relative to baseline (**Fig 1C**: paired t-test, n=10 Males: t=15.53, \*\*\*\**p*<0.0001; Females: t=6.875, \*\*\*\**p*<0.0001).

**Figure 1:**
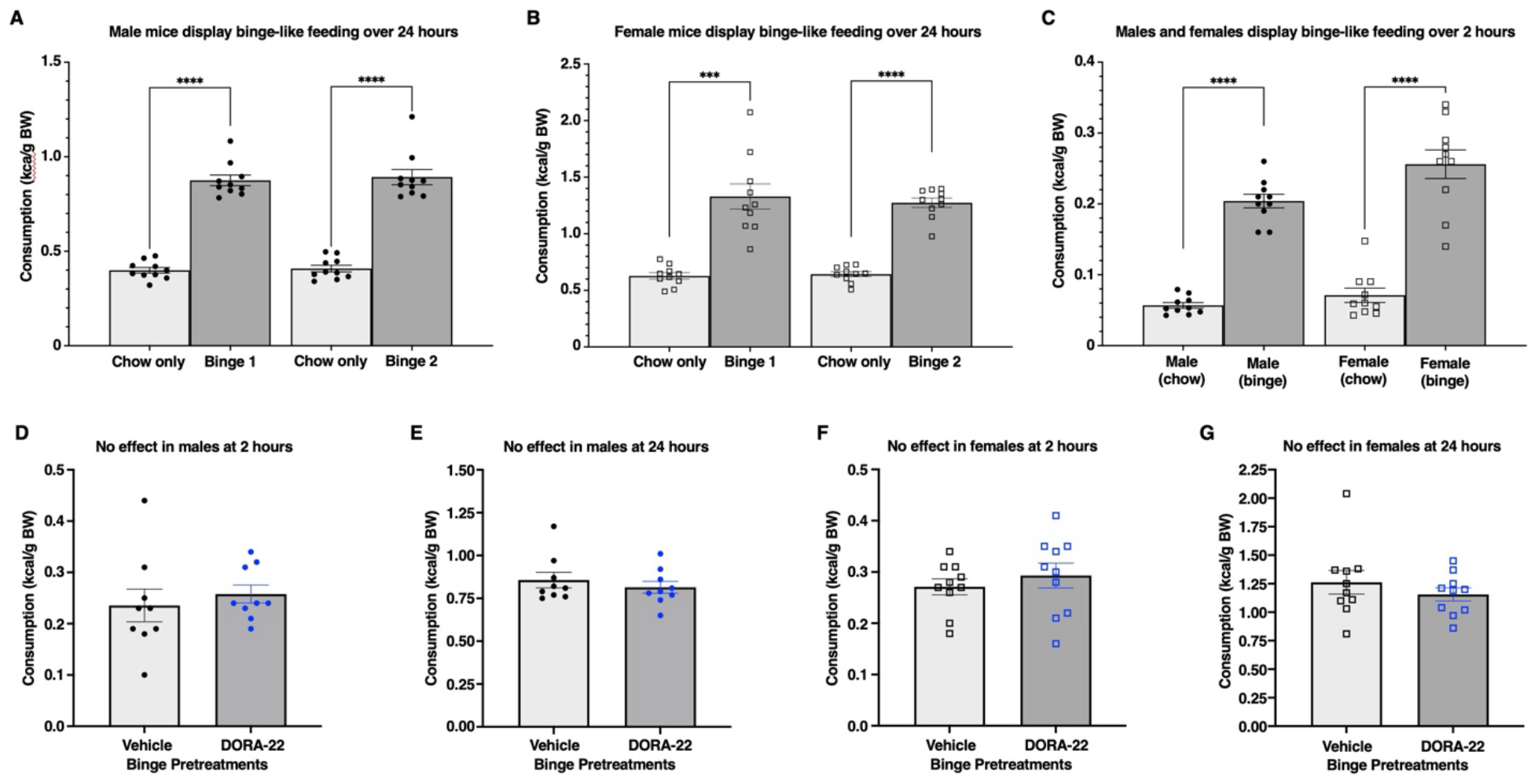
Male and female mice display binge-like behavior when intermittently exposed to a palatable diet. All C57BL/6J mice were pre-exposed to the high-fat diet for 48 hours with std. chow, and then intermittently exposed the high-fat diet (along with std. chow) for 24 hours each binge session. **1A, B)** Food intake was assessed over 24 hours for both male (n=10) and female (n=10) mice for the day prior (chow only) to a binge session and then a 24-hour binge session (chow with the high-fat diet). Males and females displayed binge-like feeding behavior with highly significant increases in total kcal/g BW consumed during binge sessions compared to the chow-only consumption. **C)** A 2-hour timepoint (to match food intake measurements after treatments) revealed significant binge-like behavior compared to a chow-only session for males and females. **D-G)** No effect of DORA-22 (100 mg/kg) was observed on binge-like feeding at the 2- or 24-hour timepoints for both sexes (n= 9 males, 10 females). All error bars are S.E.M.

### DORA-22 does not alter bingeing behavior in males or females

Mice received DORA-22 (100 mg/kg, gavage) or vehicle 20–40 minutes before the onset of the binge session, and intake was quantified at both the 2-hour and 24-hour timepoints. Across all comparisons, DORA-22 failed to reduce consumption relative to vehicle, with no detectable effect in males or females at either the early binge phase or over the full 24-hour access period (**Fig. 1D–G**). These null results were observed despite robust binge-like escalation in the same animals, indicating that dual OX1R/OX2R antagonism is insufficient, under these conditions, to attenuate this form of high-fat overeating.

### 1-SORA-51 reduces bingeing behavior in both males and females

Mice received 1-SORA-51 (100 mg/kg, gavage) or vehicle 20-40 min prior to the start of the binge sessions. In both males and females, 1-SORA-51 significantly reduced intake during the first 2 hours of high-fat access, consistent with a selective attenuation of the initial binge phase (**Fig. 2A**, males, Welch’s unpaired t-test, t=2.413, *\*p*=0.0289, n=10,8; **Fig. 2C**, females, Welch’s unpaired t-test, t=2.331, \**p*=0.0353, n=8,9). No differences were observed at 24 hours, indicating that the anorectic effect of OX1R antagonism is transient and confined to the early portion of the binge (**Fig. 2B, D**). These data contrast sharply with the null effects of the dual antagonist DORA-22, suggesting that OX1R blockade may selectively target processes that drive the initiation of binge-like consumption.

**Figure 2:**
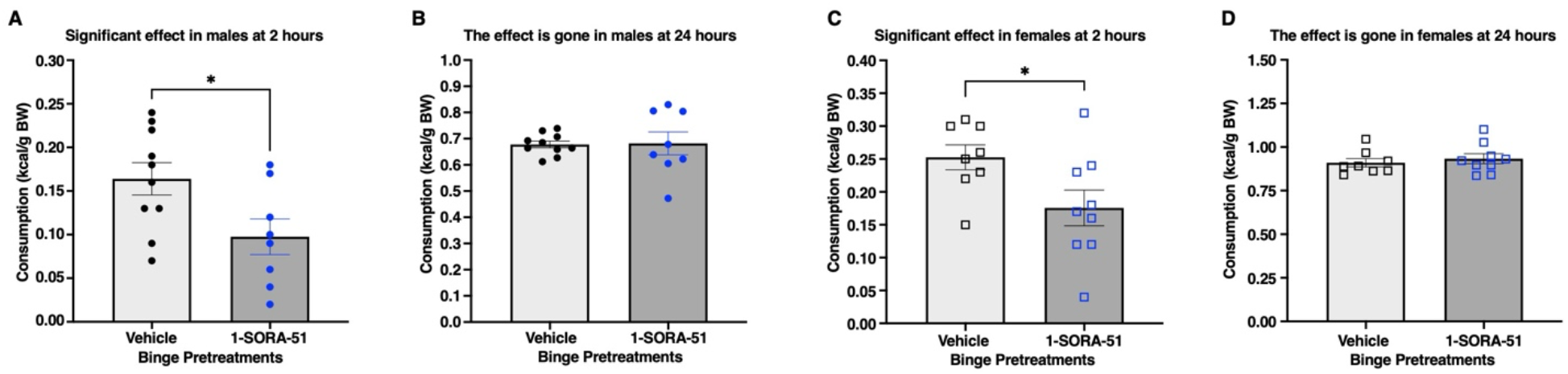
Male and female mice show a reduction in binge-like behavior after treatment with 1-SORA-51. All C57BL/6J mice were pre-exposed to the high-fat diet for 48 hours with std. chow, and then intermittently exposed the high-fat diet (along with std. chow) for 24 hours each binge session. **A)** A significant reduction in binge-like feeding was observed at the 2-hour timepoint for male mice treated with 1-SORA-51 (100 mg/kg) compared to vehicle treatment sessions (vehicle n=10, 1-SORA-51 n=8). **B)** The effect of 1-SORA-51 did not last across the 24-hour period. **C)** Similarly, a significant reduction in binge-like feeding was observed at the 2-hour timepoint for female mice treated with 1-SORA-51 (100 mg/kg) compared to vehicle treatment sessions (vehicle n=8, 1-SORA-51 n=9). **D)** The effect of 1-SORA-51 did not last across the 24-hour period. All error bars are S.E.M.

### 1-SORA-51 reduces bingeing behavior in a dopamine-independent manner

To determine whether the behavioral effects of OX1R antagonism were accompanied by alterations in mesolimbic dopamine signaling, we recorded nucleus accumbens GRABDA_2m_ fluorescence during binge sessions in mice undergoing counterbalanced vehicle and 1-SORA-51 treatments. Male and female mice expressing the dopamine GRABDA_2m_ sensor in the nucleus accumbens were subjected to the binge training paradigm. On binge days, fiber photometry recordings were taken during habituation, gavage, and high-fat pellet exposure. Sessions were analyzed in 2-min bins over 30 minutes after high-fat pellet exposure, and 10-min bins over 120 minutes after high-fat pellet exposure. Across both sexes, baseline fluorescence, post-gavage-associated activity, and the characteristic dopamine increase elicited by high-fat pellet delivery were indistinguishable between drug and vehicle conditions, indicating that 1-SORA-51 does not measurably alter dopamine dynamics before or during the binge (**Fig. 3A**). Notably, both sessions revealed a similar increase in fluorescence upon high-fat pellet delivery. Despite this neurochemical similarity, 1-SORA-51 produced a significant reduction in high-fat consumption within the same animals, replicating its behavioral effects in independent cohorts (**Fig 3B**: paired t-test, t=2.675, \**p*=0.0281, n=9). These findings demonstrate that OX1R antagonism suppresses early binge-like intake without detectable modulation of nucleus accumbens dopamine release, suggesting that its anorectic action engages dopamine-independent motivational processes.

**Figure 3:**
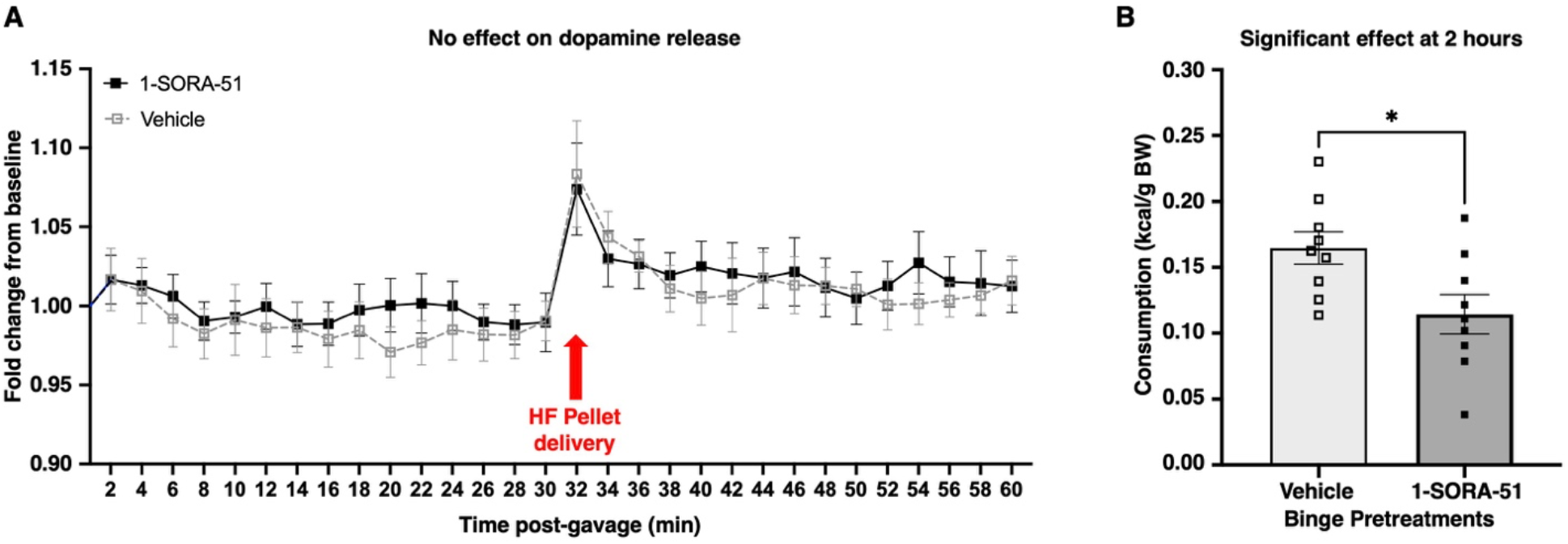
Treatment with 1-SORA-51 reduces binge-like behavior without altering dopamine release in the nucleus accumbens. **A)** Male and female mice (n=9) expressing the GRABDA_2m_ sensor and finished binge training underwent fiber photometry sessions, during which baseline was recorded prior to treatment via gavage, and then provided access to the high-fat diet for 2 hours. Shown here is fluorescence normalized to baseline. No effect of treatment was seen during the 30 minutes after gavage or after HF pellet delivery. **B)** Treatment with 1-SORA-51 resulted in significantly less high-fat pellet consumption compared to vehicle sessions. All error bars are S.E.M.

## Methods

### Sex as a biological variable

Male and female mice were used for these studies. No sex-specific effects were found upon analysis.

### Animals

Animal experiments were carried out in accordance with Yale University School of Medicine Institutional Animal Care and Use Committee (IACUC) regulations. All protocols were approved by the Yale IACUC. Male and female C57BL/6J mice, ages 8-12 weeks at the start of these studies were used (Jackson Laboratories, Bar Harbor). All methods are presented in accordance with ARRIVE guidelines with the exception that the experimenters were not blinded to the treatments. Animals that experienced difficult gavage attempts, or whose eating behavior (occasional shredding) made food intake measurements impossible were removed, and these included: 1) One male mouse from the behavioral DORA-22 study (for vehicle and DORA-22), 2) Two males from the behavioral 1-SORA-51 treatment group, and 3) Two females from the vehicle and one female from the behavioral 1-SORA-51 treatment group. Mice were group housed up to five per cage and maintained *ad libitum* with standard chow (#**2918** Teklad Global diet, Inotiv Inc.) on a 12-hour light/dark cycle (lights on at 7 a.m.), and then single-housed prior to starting the bingeing experiments. For fiber photometry studies, pAAV-hsyn-GRAB-DA2m with a titer of 2.4×10^13^ GC/ml was purchased from Addgene (Watertown, MA). Mice were anesthetized with isoflurane, treated with analgesics, and underwent intracranial viral infusion with GRAB-DA2m, delivering 0.5 µl unilaterally to the right nucleus accumbens (stereotaxic coordinates A/P +1.2, M/L +0.6, D/V -4.5 mm from bregma) over five minutes, with an additional five minutes to allow for diffusion before retracting the Hamilton syringe. A fiber optic cannula (400 µm dia. fiber core, N.A. 0.48) was lowered to the same A/P and M/L coordinates, and D/V -4.2 mm (from bregma), and then cemented in place. Postoperative analgesics were administered, and experimentation began no sooner than two weeks later to allow for mice to fully recover and for adequate viral expression to occur.

### Bingeing paradigm

Mice were initially exposed to a “western” high-fat diet for 48 hours while group housed and with standard chow included as well. The high-fat diet used for all studies (5TJN, TestDiet, Richmond, Indiana) provides 4.55 kcal/g with the following caloric breakdown: protein 15.8%, fat 39.1%, and carbohydrates 45.1%. For behavioral bingeing studies, mice were single-housed after their initial exposure to the high-fat diet. All mice entered the cycle of 6 days of *ad libitum* maintenance with standard chow alone followed by one day of bingeing with high-fat diet (just prior to the start of the dark cycle) along with standard chow. Cages and bedding were changed after each binge session to remove any high-fat diet remnants. This cycle was repeated throughout the experiments. For fiber photometry studies, mice remained group housed throughout the initial exposure and intermittent testing sessions, which were conducted during the light cycle.

### Behavioral Binge Pharmacology

DORA-22 and 1-SORA-51 were provided by Merck, Inc. The vehicle used to deliver these compounds via gavage was 10% Tween 80. The dose of 100 mg/kg for both exceeded solubility limits, so mechanical grinding with a pestle attached to an electric drill was required to reduce the size of the solids to allow for passage through the gavage tip. Mixing was necessary to homogenize the particle suspension just prior to gavaging each mouse. Treatments occurred approximately 20-40 minutes prior to the start of the dark cycle. Chow and high-fat pellets were measured and placed on the floor of their cage just prior to the start of the dark cycle and measured again 2 hours into the binge session. Body weight and all foods were assessed at 24 hours, with care to locate measurable remnants in the bedding, after which all cages were changed. Testing involved delivering either vehicle or compound and then reversing the treatments in the same animals a week later for the next binge session test.

### Fiber Photometry

The fiber photometry system consisted of a TDT RZ5 module (Tucker-Davis Technologies, Alachua, FL), an LED driver with a 490 nm cube (all from Thorlabs, Newton, NJ), and a photodetector (Newport #2151, Irvine, CA). The emission spectrum passed through a 525 ± 39 nm bandpass filter prior to photodetection, and these data were sampled by the TDT RZ5 module at a rate of 381.4697 Hz. Mice remained group housed after surgery, through binge training and testing. After viral expression, mice were habituated to the testing room and the fiber optic to reduce any novelty effects prior to experimentation. Test sessions consisted of initially photobleaching the fiber optic tether for at least 30 minutes at the beginning of each day of testing, prior to attaching a mouse. Mice were then tethered to a fiber optic cable (400 µm dia. core fiber, N.A. 0.48, Doric Lenses, Quebec, Canada) and placed in a plastic mouse bucket with home cage bedding for 30 minutes for general habituation with the LED turned off. Buckets and bedding were replaced when switching between cages of mice. After habituation, the LED was turned on and 1 hour of baseline fluorescence was recorded before gavage. 30 minutes after gavage, a high-fat pellet was placed in the bedding, and the recording continued for 2 hours. All mice went through vehicle and 1-SORA-51 treatment sessions on their binge days.

### MATLAB analysis

Data for the fiber photometry setup were collected as intensity from recorded filtered emission spectrum generated by the 490 nm wavelength. Recordings were continuous throughout the session, and fluorescent data recorded while investigators were in the room or handling the mice were removed from the data set prior to any normalization and baseline calculation. Handling typically did not exceed one minute. After removing the signal during handling/treatment, the average signal during the pre-drug baseline was calculated and used to compute fold-change in fluorescence for the entire session. Area under the curve was then calculated for this fold change in two-minute bins. This method of using just a single signal for analysis without a valid isosbestic control is similar to that used by others for GRAB sensors (Trinko et al. 2024; Basu et al. 2024; Feng et al. 2024; Li et al. 2023).

### Statistics

GraphPad Prism 10 was used for statistical analysis of fold change values. Two-tailed t-tests were used for food intake comparisons. In the event of excluding a mouse due to the inability to accurately measure food intake, a Welch’s unpaired t-test was performed. All other t-tests were paired. Two-way RM-ANOVAs were used for comparing the fiber photometry signal between treatments. An alpha threshold of 0.05 was used for statistical significance.

### Study approval

All animal care and experimental procedures were conducted in accordance with the Yale University School of Medicine Institutional Animal Care and Use Committee (IACUC) guidelines.

## Data availability

All MATLAB code will be made accessible upon request through the corresponding author.

## Discussion

The present study demonstrates that selective OX1R antagonism, but not dual OX1R/OX2R blockade, suppresses the initiation of binge-like feeding in a non-stress-based intermittent high-fat paradigm. Although DORA-22 and 1-SORA-51 were administered under identical conditions, only OX1R antagonism reduced consumption, and this effect was confined to the first two hours of access. Fiber photometry revealed similar nucleus accumbens dopamine dynamics across drug and vehicle sessions, indicating that the behavioral effect occurred without detectable modulation of mesolimbic dopamine release. These findings identify OX1R signaling as a critical driver of the early motivational phase of binge-like intake.

The efficacy of 1-SORA-51 aligns with established roles of OX1R pathways in regulating reward seeking, cue reactivity, and the vigor of approach behavior. OX1R-expressing targets in mesolimbic circuits receive dense orexin input, and their engagement is thought to amplify behavioral activation in response to salient cues. The selective reduction in early—but not 24-hour—intake suggests that OX1R may contribute preferentially to the processes governing binge initiation, rather than to longer-term caloric regulation. The use of a nondeprived, palatability-driven model strengthens this interpretation by eliminating restriction-induced homeostatic feeding influences. However, the relatively short half-life of 1-SORA-51 (3-4 hours) may also contribute to the lack of effect at the longer time point.

In contrast, the lack of effect of DORA-22 highlights the functional divergence between OX1R and OX2R pathways. Given OX2R’s established role in sleep–wake stabilization, dual antagonism may compromise arousal without strongly impacting motivational circuits relevant to binge initiation, or OX2R blockade may mask behavioral effects attributable to OX1R inhibition. These findings underscore the importance of receptor specificity when evaluating orexin-directed pharmacology and suggest that OX1R-selective compounds may offer a more targeted approach for modulating maladaptive consumption.

The photometry data further refine the mechanistic interpretation. High-fat pellet delivery elicited the expected phasic dopamine response, and neither its amplitude nor its temporal profile was altered by 1-SORA-51. This is consistent with microdialysis data indicating no detectable effect of intra-accumbens OX1R blockade with SB-334867 on basal accumbal dopamine efflux (Kawashima et al. 2022). Thus, the attenuation of binge-like intake is unlikely to reflect a gross blunting of phasic dopamine responses to food delivery. Instead, the effect might arise upstream or parallel to dopamine release, potentially within circuits encoding arousal, salience, or behavioral mobilization. This supports models in which orexin contributes to motivational drive without necessarily modifying stimulus-evoked dopamine transients.

Methodological heterogeneity across binge-eating paradigms complicates comparisons in the literature, as stress-based, deprivation-based, or limited-access models engage distinct physiological states. By using the intermittent high-fat paradigm, which isolates opportunity-and palatability-driven intake (Czyzyk et al. 2010), the present study situates the behavioral effects of OX1R antagonism during highly motivated behavior. Under these conditions, OX1R blockade selectively targets the rapid escalation characteristic of binge onset—a feature with strong relevance for binge eating disorder and related conditions.

Translationally, these findings support OX1R-selective antagonism as a potential strategy for reducing binge propensity without directly suppressing mesolimbic dopamine signaling. The divergence between DORA-22 and 1-SORA-51 emphasizes that receptor-level specificity matters for therapeutic design and cautions against assuming that compounds developed for insomnia will generalize uniformly to feeding behavior. Overall, these results clarify how orexin signaling contributes to maladaptive high-fat consumption and provide a framework for exploring OX1R-targeted approaches in binge eating and other disorders of dysregulated approach behavior.

## References

Alcaraz-Iborra, Manuel, Francisca Carvajal, José Manuel Lerma-Cabrera, Luis Miguel Valor, and Inmaculada Cubero. 2014. “Binge-like Consumption of Caloric and Non-Caloric Palatable Substances in Ad Libitum-Fed C57BL/6J Mice: Pharmacological and Molecular Evidence of Orexin Involvement.” Behavioural Brain Research 272 (October): 93–99. 10.1016/jLbbr.2014.06.049.

Basu, Aakash, Jen-Hau Yang, Abigail Yu, et al. 2024. “Frontal Norepinephrine Represents a Threat Prediction Error Under Uncertainty.” Biological Psychiatry, Monoaminergic Dysregulation in Posttraumatic Stress Disorder—Revisited, vol. 96 (4): 256-67. 10.1016/j.biopsych.2024.01.025.

Brown, Robyn Mary, Andrezza K. Kim, Shaun Yon-Seng Khoo, Jee Hyun Kim, Bianca Jupp, and Andrew John Lawrence. 2016. “Orexin-1 Receptor Signalling in the Prelimbic Cortex and Ventral Tegmental Area Regulates Cue-Induced Reinstatement of Ethanol-Seeking in iP Rats.” Addiction Biology 21 (3): 603–12. 10.1111/adb.12251.

Cai, X J, P S Widdowson, J Harrold, et al. 1999. “Hypothalamic Orexin Expression: Modulation by Blood Glucose and Feeding.” Diabetes 48 (11): 2132–37. 10.2337/diabetes.48.11.2132.

Carr, Kenneth D. 2002. “Augmentation of Drug Reward by Chronic Food Restriction: Behavioral Evidence and Underlying Mechanisms.” Physiology & Behavior, Proceedings from the 2001 Meeting of the Society for the Study of Ingestive Behavior (SSIB), vol. 76 (3): 35364. 10.1016/S0031-9384(02)00759-X.

Cluderay, J. E, D. C Harrison, and G. J Hervieu. 2002. “Protein Distribution of the Orexin-2 Receptor in the Rat Central Nervous System.” Regulatory Peptides, Special Issue: Orexin, vol. 104 (1): 131–44. 10.1016/S0167-0115(01)00357-3.

Corwin, Rebecca L, and Ariel Buda-Levin. 2004. “Behavioral Models of Binge-Type Eating.” Physiology & Behavior, Festschrift in Honor of Gerard P. Smith, vol. 82 (1): 123–30. 10.1016/j.physbeh.2004.04.036.

Cowin, Rebecca L, Nicole M. Avena, and Mary M. Boggiano. 2011. “Feeding and Reward: Perspectives from Three Rat Models of Binge Eating.” Physiology & Behavior 104 (1): 8797. 10.1016/j.physbeh.2011.04.041.

Czyzyk, Traci A., Allison E. Sahr, and Michael A. Statnick. 2010. “A Model of Binge-Like Eating Behavior in Mice That Does Not Require Food Deprivation or Stress.” Obesity 18 (9): 1710–17. 10.1038/oby.2010.46.

Diagnostic and Statistical Manual of Mental Disorders, Fifth Edition. 2013. 5th ed. American Psychiatric Association.

Feng, Jiesi, Hui Dong, Julieta E. Lischinsky, et al. 2024. “Monitoring Norepinephrine Release in Vivo Using next-Generation GRABNE Sensors.” Neuron 112 (12): 1930-1942.e6. 10.10167j.neuron.2024.03.001.

Hardaway, J. A., N. A. Crowley, C. M. Bulik, and T. L. Kash. 2015. “Integrated Circuits and Molecular Components for Stress and Feeding: Implications for Eating Disorders.” Genes, Brain, and Behavior 14 (1): 85–97. 10.1111/gbb.12185.

Herring, W. Joseph, Kathryn M. Connor, Neely Ivgy-May, et al. 2016. “Suvorexant in Patients With Insomnia: Results From Two 3-Month Randomized Controlled Clinical Trials.” Biological Psychiatry, Borderline Personality Disorder: Mechanisms of Emotion Dysregulation, vol. 79 (2): 136–48. 10.1016/j.biopsych.2014.10.003.

Hudson, James I., Eva Hiripi, Harrison G. Pope, and Ronald C. Kessler. 2007. “The Prevalence and Correlates of Eating Disorders in the National Comorbidity Survey Replication.” Biological Psychiatry 61 (3): 348–58. 10.1016Zj.biopsych.2006.03.040.

Kawashima, Hiroki, Yuri Aono, Yuriko Watanabe, John L. Waddington, and Tadashi Saigusa. 2022. “In Vivo Microdialysis Reveals That Blockade of Accumbal Orexin OX2 but Not OX1 Receptors Enhances Dopamine Efflux in the Nucleus Accumbens of Freely Moving Rats.” The European Journal of Neuroscience 55 (3): 733–45. 10.1111/ejn.15593.

Lai, Francesco, Flavia Cucca, Roberto Frau, et al. 2018. “Systemic Administration of Orexin a Loaded Liposomes Potentiates Nucleus Accumbens Shell Dopamine Release by Sucrose Feeding.” Frontiers in Psychiatry 9 (December): 640. 10.3389/fpsyt.2018.00640.

Lecea, L. de, T. S. Kilduff, C. Peyron, et al. 1998. “The Hypocretins: Hypothalamus-Specific Peptides with Neuroexcitatory Activity.” Proceedings of the National Academy of Sciences of the United States of America 95 (1): 322–27. 10.1073/pnas.95.1.322.

Levin, Barry E. 2006. “Orexins: Neuropeptides for All Seasons and Functions.” American Journal of Physiology-Regulatory, Integrative and Comparative Physiology 291 (4): R885–88. 10.1152/ajpregu.00344.2006.

Li, Li, Akshay N. Rana, Esther M. Li, Jiesi Feng, Yulong Li, and Michael R. Bruchas. 2023. “ActivityDependent Constraints on Catecholamine Signaling.” Cell Reports 42 (12): 113566. 10.1016/j.celrep.2023.113566.

Lin, Ling, Juliette Faraco, Robin Li, et al. 1999. “The Sleep Disorder Canine Narcolepsy Is Caused by a Mutation in the Hypocretin (Orexin) Receptor 2 Gene.” Cell 98 (3): 365–76. 10.1016/S0092-8674(00)81965-0.

Marcus, J. N., C. J. Aschkenasi, C. E. Lee, et al. 2001. “Differential Expression of Orexin Receptors 1 and 2 in the Rat Brain.” The Journal of Comparative Neurology 435 (1): 6–25. 10.1002/cne.1190.

Piccoli, Laura, Maria Vittoria Micioni Di Bonaventura, Carlo Cifani, et al. 2012. “Role of Orexin-1 Receptor Mechanisms on Compulsive Food Consumption in a Model of Binge Eating in Female Rats.” Neuropsychopharmacology: Official Publication of the American College of Neuropsychopharmacology 37 (9): 1999–2011. 10.1038/npp.2012.48.

Rehn, Simone, Joel S. Raymond, Robert A. Boakes, Michael D. Kendig, and Cathalijn H. C. Leenaars. 2025. “Behavioural and Physiological Effects of Binge Eating: A Systematic Review and Meta-Analysis of Animal Models.” Neuroscience & Biobehavioral Reviews 173 (June): 106135. 10.1016/jmeubiorev.2025.106135.

Rossi, Mark A., and Garret D. Stuber. 2018. “Overlapping Brain Circuits for Homeostatic and Hedonic Feeding.” Cell Metabolism 27 (1): 42–56. 10.1016Zj.cmet.2017.09.021.

Sakurai, T., A. Amemiya, M. Ishii, et al. 1998. “Orexins and Orexin Receptors: A Family of Hypothalamic Neuropeptides and G Protein-Coupled Receptors That Regulate Feeding Behavior.” Cell 92 (4): 573–85. 10.1016/s0092-8674(00)80949-6.

Swanson, Sonja A., Scott J. Crow, Daniel Le Grange, Joel Swendsen, and Kathleen R. Merikangas. 2011. “Prevalence and Correlates of Eating Disorders in Adolescents. Results from the National Comorbidity Survey Replication Adolescent Supplement.” Archives of General Psychiatry 68 (7): 714–23. 10.1001/archgenpsychiatry.2011.22.

Thorpe, A. J., and C. M. Kotz. 2005. “Orexin A in the Nucleus Accumbens Stimulates Feeding and Locomotor Activity.” Brain Research 1050 (1): 156–62. 10.1016/j.brainres.2005.05.045.

Tomasi, Dardo, and Nora D. Volkow. 2013. “Striatocortical Pathway Dysfunction in Addiction and Obesity: Differences and Similarities.” Critical Reviews in Biochemistry and Molecular Biology 48 (1): 1–19. 10.3109/10409238.2012.735642.

Trinko, Joseph R., Ethan Foscue, Edward M. Kong, et al. 2024. “Xylazine Induces Dopamine Release and Augments the Effects of Fentanyl.” The Journal of Clinical Investigation 134 (22). 10.1172/JCI183354.

Udo, Tomoko, and Carlos M. Grilo. 2018. “Prevalence and Correlates of DSM-5-Defined Eating Disorders in a Nationally Representative Sample of U.S. Adults.” Biological Psychiatry, Habits and Compulsion, vol. 84 (5): 345–54. 10.1016/j.biopsych.2018.03.014.

Volkow, N. D., G.-J. Wang, D. Tomasi, and R. D. Baler. 2013. “Pro v Con Reviews: Is Food Addictive?” Obesity Reviews : An Official Journal of the International Association for the Study of Obesity 14 (1): 2–18. 10.1111/j.1467-789X.2012.01031.x.

Wang, Gene-Jack, Allan Geliebter, Nora D. Volkow, et al. 2011. “Enhanced Striatal Dopamine Release During Food Stimulation in Binge Eating Disorder.” Obesity (Silver Spring, Md.) 19 (8): 1601–8. 10.1038/oby.2011.27.

Winrow, Christopher J., Anthony L. Gotter, Christopher D. Cox, et al. 2011. “Promotion of Sleep by Suvorexant—A Novel Dual Orexin Receptor Antagonist.” Journal of Neurogenetics 25 (1-2): 52–61. 10.3109/01677063.2011.566953.

Yu, Yang, Renee Miller, and Susan W. Groth. 2022. “A Literature Review of Dopamine in Binge Eating.” Journal of Eating Disorders 10 (1): 11. 10.1186/s40337-022-00531-y.

